# Contrastive Fitness Learning: Reprogramming Protein Language Models for Low-*N* Learning of Protein Fitness Landscape

**DOI:** 10.1101/2024.02.11.579859

**Authors:** Junming Zhao, Chao Zhang, Yunan Luo

**Affiliations:** School of Computational Science and Engineering, Georgia Institute of Technology

## Abstract

Machine learning (ML) is revolutionizing our ability to model the fitness landscape of protein sequences, which is critical to answering fundamental life science questions and addressing important protein engineering applications, such as quantifying the pathogenicity of disease variants, forecasting viral evolution in a pandemic, and engineering new antibodies. Recently, the protein language model (pLM) has emerged as an effective ML tool in deciphering the intrinsic semantics of protein sequences and become the foundation of state-of-the-art ML solutions for many problems in protein biology. However, significant challenges remain in leveraging pLMs for protein fitness prediction, in part due to the disparity between the scarce number of sequences functionally characterized by high-throughput assays and the massive data samples required for training large pLMs. To bridge this gap, we introduce Contrastive Fitness Learning (ConFit), a pLM-based ML method for learning the protein fitness landscape with limited experimental fitness measurements as training data. We propose a novel contrastive learning strategy to fine-tune the pre-trained pLM, tailoring it to achieve protein-specific fitness prediction while avoiding overfitting, even when using a small number (low-*N*) of functionally assayed mutant sequences for supervised fine-tuning. Evaluated across over 30 benchmark datasets of protein fitness, ConFit consistently provided accurate fitness predictions and outperformed several competitive baseline methods. Further analysis revealed that ConFit’s capability of low-*N* learning enabled sample-efficient active learning for identifying high-fitness protein variants. Collectively, our work represents a novel strategy to harness the potential of pLMs to elucidate the protein sequence-function relationship. The source code of ConFit is available at https://github.com/luo-group/ConFit.

## 1 Introduction

Evolution has shaped natural proteins to perform vital functions in life science and address pressing applications ranging from developing targeted therapeutics ^1^ to engineering herbicide-resistant crops ^2^ and sustainable biocatalysis ^3^. The function capabilities of naturally occurring proteins are further expanded through a biotechnology technique known as protein engineering, which enhances a natural protein’s original function (e.g., improving an enzyme’s catalytic activity ^4^) or repurposes it to a different but related function (e.g., repurposing an antibody to target a new virus ^5^), often by introducing “beneficial” mutations into the protein’s amino acid sequence. A protein’s function is encoded by its amino acid sequence, and the sequence-fitness landscape maps out the relationships between a protein sequence and function. The goal of protein engineering is to search this landscape for high-fitness sequences. Here. ‘fitness’ broadly refers to protein properties such as ligand-binding affinity, stability, or catalytic activity. Despite the success in practice ^6;7^, using protein engineering to design or discover new novel proteins with desired functions is still challenging because i) the laboratory experiments are time- and resource-intensive and ii) the sequence search space is astronomically large (scaling exponentially with protein length).

Machine learning (ML) has emerged as an effective approach to infer the sequence-fitness relationship. Used as surrogate models, ML methods, including both supervised and unsupervised models, predict the properties of proteins more rapidly and less expensively than wet-lab experiments, reducing the overall experimental burden in protein engineering. Unsupervised generative ML models have been developed to learn the intrinsic semantics in naturally occurring protein sequences. Although not labeled with measurements for the property of interest, those sequences are subjected to evolutionary pressure and implicitly encode sequence constraints that are prerequisites for protein fitness. To capture those constraints, unsupervised approaches often fit a probability density model (e.g., hidden Markov models ^8^, Potts models ^9–19^, latent variable models ^20–22^, and protein language models ^23–29^) on natural sequences – either homologous sequences of the target protein or massive natural protein sequences from databases such as UniProt ^30^ – to estimate the occurrence probability *p*(***x***) of a particular sequence ***x*** as a proxy of its fitness ^9^.

Supervised models are particularly useful for explicitly learning the sequence-fitness relationship when paired data of variant sequences and experimental fitness are richly available, often outperforming unsupervised models ^31–34^. Yet, their effectiveness is constrained by the scarcity of fitness data. Screening variant fitness can be lab-intensive and require developing tailored assays for a specific function, with typical experiments yielding only 10^0^-10^3^ labeled variants. However, modern ML models such as neural networks are notoriously data-hungry and often require *>*10^5^ training samples. While high-throughput assays can screen fitness at scale, they may compromise fidelity for throughput ^35^, and not all proteins have high-throughput assays. This gap has spurred the development of ML models that can make accurate fitness predictions even with small-size (‘low *N* ‘) training data. A common strategy is to leverage unsupervised models to capture general sequence patterns across natural proteins, which hold implications for functions, and then adapt those models using low-*N* fitness data to make supervised, protein-specific fitness prediction ^36–38^.

Protein language models (pLMs), in particular, are effective unsupervised models used in existing low-*N* fitness prediction methods ^36–38^ for learning sequence semantics that are likely to occur in natural proteins. Interestingly, even without fitness-labeled data, pLMs have shown accurate strong performance in predicting the effects of amino acid substitutions in protein sequences, correlating well with experimentally measured fitness data ^39–43^. Given their promising predictive capacity, we reason that with even a small number of fitness data, pLMs could be fine-tuned into more accurate supervised models. However, directly fine-tuning a pLM (typically with ∼10^9^ parameters) on sparse low-*N* data (∼10^2^ samples) will likely lead to overfitting, abruptly distorting its learned sequence patterns during pre-training – a phenomenon known as *catastrophic forgetting* in ML ^44^. Despite its prevalence in ML-guided protein engineering, only a few studies attempted to tackle it, often compromising prediction accuracy to avoid overfitting ^31;36^.

Here, we introduce ConFit (<u>Con</u>trastive <u>Fit</u>ness Learning), an ML algorithm that effectively reprograms a pre-trained pLM to high-accuracy, sample-efficient fitness prediction model under low-*N* settings. Our method represents a new paradigm of pLM fine-tuning, which preserves the evolutionary patterns the pLM has learned during pre-training while effectively incorporating small-size fitness data to enable specific and accurate fitness predictions without overfitting. Recognizing that in ML-guided protein engineering (MLPE), ranking variants by their fitness is more crucial than predicting exact fitness values for individual variants, we shift from traditional regression ^36;37^ to a contrastive learning approach. Our method calibrates the pLM to predict the *relative fitness order* between pairwise variants rather than their *absolute fitness values*. This not only expands the effective training set quadratically but also circumvents catastrophic forgetting effectively.

Benchmarking on over 30 mutagenesis datasets of protein fitness, we showed that ConFit can accurately predict protein fitness even with only *N* =48 fitness-labeled sequences as training data, outperforming other unsupervised and supervised models. The outstanding low-*N* prediction capability of our method also enabled a sample-efficient active learning loop for identifying function-enhanced variants. We expect ConFit to be a practically useful tool for guiding and accelerating protein engineering, especially for applications where experimental fitness screening is costly.

## 2 Protein Language Models for Fitness Prediction: Revisited

The advent of ChatGPT and subsequent LLMs ^45–48^ has not only revolutionized natural language processing but has also had a transformative impact on computational biology through the development of protein language models (pLMs) ^23–29^. pLMs adopt LLM strategies, applying self-supervised learning to predict masked amino acids in protein sequences given other residues as context, uncovering patterns indicative of natural protein grammar. Many state-of-the-art ML algorithms for protein biology now rely on pLMs, from the prediction of protein structure ^28^ to protein properties like solubility ^49^, stability ^24^, and binding affinity ^43^, with particular effectiveness in protein fitness prediction ^27;34;43^. The success of pLMs hinges on their extensive model size and the massive training data, allowing for learning general patterns across natural proteins that transfer well to other problems where labeled data are limited.

Formally, we denote the amino acid (AA) sequence of a protein by ***x*** = (*x*_1_, …, *x*_*L*_) ∈ *X* ^*L*^, where *x*_*i*_ is the *i*-th AA, *L* the sequence length, and *X* the alphabet of possible AAs (e.g., the 20 standard AAs). Masked pLMs ^27;28^ aims to learn the conditional probability *p*(*x*_*i*_|*x*_−*i*_) of an AA appearing at a given position, conditioned on the sequence excluding that position, where *x*_−*i*_ = (*x*_1_, …, *x*_*i*−1_, *x*_*i*+1_, …, *x*_*L*_) (Fig 1a). While our focus is on masked pLMs in this work, we acknowledge alternative models like autoregressive pLMs ^29;50^, which estimate *p*(*x*_*i*_|*x*_*<i*_), taking into account all preceding AAs *x*_*<i*_ = (*x*_1_, …, *x*_*i*−1_).

**Figure 1:**
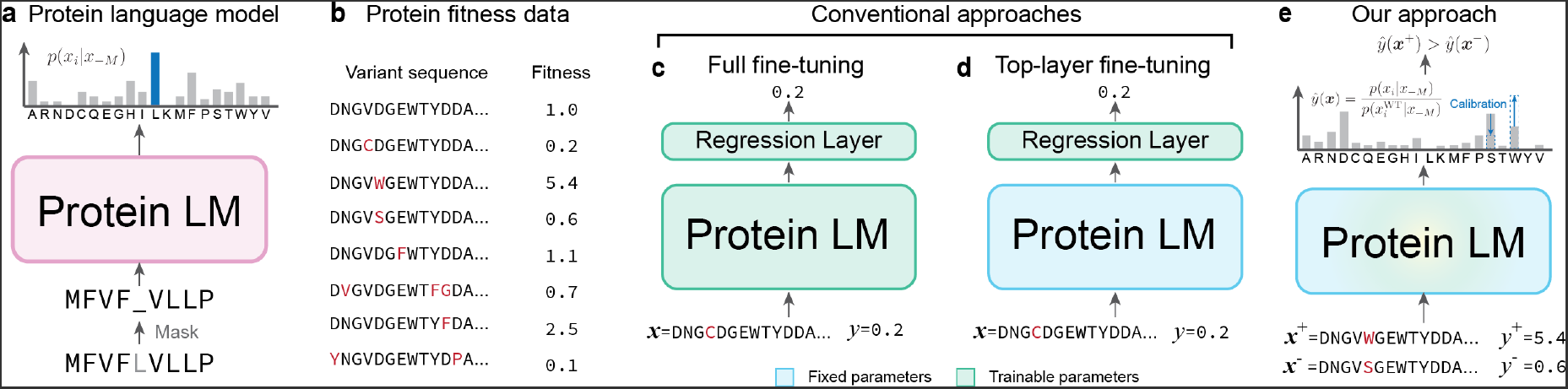
Overview of ConFit. (a) A protein language model (pLM) predicts the probability of amino acid at position given other unmasked positions. (b) Example of protein fitness data in which the variants of a wildtype (WT) sequence are experimentally characterized with fitness values. (c-d) Conventional approaches for fine-tuning pre-trained pLMs for protein fitness prediction, including “full fine-tuning” which updates all parameters in the pLM, and “top-layer fine-tuning” which trains the top-layer on the fitness data while fixing other pre-trained parameters. (e) Our approach, ConFit, calibrates pre-trained pLM for low-*N* fitness prediction through contrastive learning.

The natural protein sequences observed today are the culmination of evolutionary selection for fitness and function. Trained on massive natural protein sequence database such as UniProt ^30^, pLMs implicitly capture the sequence patterns that are important for fitness. Specifically, the conditional probability *p*(*x*_*i*_|*x*_−*i*_) estimated by pLMs gives a probability distribution over *X* for a given site *i*, indicating the site’s ‘evolutionary preference’ for an AA *x*_*i*_ ∈ *X* . A higher *p*(*x*_*i*_|*x*_−*i*_) intimates an AA *x*_*i*_ that is evolutionarily advantageous, presumably contributing positively to fitness, while a lower value suggests potentially deleterious mutations.

Following this rationale, recent studies directly used the PLM-predicted likelihood for zero-shot predictions of AA mutation effects. Even without any supervision from paired sequence-fitness data, pLM’s predictions correlate strongly with laboratory-measured protein fitness (e.g., deep mutagenesis scans) ^39–41^. PLMs have thus been harnessed for important biomedical applications, including recommendation mutations to enhance antibody binding ^43^ or predicting the pathogenicity of human missense variants ^51^.

Given that the pLMs have strong zero-shot fitness prediction performance despite having no prior exposure to fitness data, one may reason that augmenting such models with even a modest number of functionally characterized variants as training data (Fig 1b) could significantly improve their fitness prediction accuracy. The intuitive strategy might be to fine-tune the pLM with available fitness data (Fig 1c), following common practice in ML ^52^. However, the unique challenge in protein engineering is that current quantitative assays to measure fitness are not scalable to high throughput, yielding only tens to hundreds of fitness measurements. This paucity of data, juxtaposed against the millions or even billions of parameters in pLMs, collectively presents a challenge where directly fine-tuning pLMs on such sparse data easily leads to catastrophic forgetting, distorting the pLM’s pre-trained protein sequence features and causing the model to overfit the sparse fitness data.

To mitigate catastrophic forgetting in low-*N* fitness prediction, some studies have adopted a compromised approach: freezing the pLM parameters while introducing a new trainable regression layer—typically a ridge or LASSO model—on top of pLM-generated sequence representations ^36;37^ (Fig 1d). This hybrid model, with significantly fewer trainable parameters, is less prone to overfitting and has been shown to accurately predict fitness even using only *N* = 24 or 48 variants as training data ^36;37^. However, these approaches mitigated catastrophic forgetting at the expense of limited prediction accuracy: freezing the pLM’s layers forfeits its capacity to capture complex epistatic interactions ^53;54^ inherent in protein sequences—a task linear models find challenging. To date, how to fully harness the pre-learned knowledge in pLMs for low-*N* fitness prediction, without catastrophic forgetting and overfitting, remains an unresolved challenge.

## 3 Methods

We present ConFit, a pLM-based ML method for low-*N* protein fitness prediction. Our algorithm is motivated by a key observation of a mismatch exists between the pre-training and fine-tuning learning objectives, which leads to catastrophic forgetting during the latter. We propose a principled way to achieve robust low*N* pLM fine-tuning for protein fitness prediction. Different from existing methods that either fully fine-tune the pLM (“full fine-tuning”), suffering from severe overfitting in low-*N* scenarios, or only re-fit the pLM’s top layer while keeping the rest untouched (“top-layer fine-tuning”), resulting in compromised accuracy, our method provides both high accuracy and high sample efficiency for low-*N* protein engineering ^36^.

### 3.1 Motivation and Rationale of ConFit

The motivation behind ConFit lies in the realization that catastrophic forgetting occurs when a pLM is fine-tuned for a task divergent from its initial training. Originally, pLMs maximized the likelihood of natural protein sequences sampled from the training data, i.e., max 𝔼_***x*∏***i*_ *p*(*x*_*i*_|*x*_−*i*_). However, during fine-tuning for fitness prediction, the objective shifts to minimizing the difference between observed and predicted fitness values, measured by mean squared error (MSE): min Σ_*k*_ ∥*y*^(*k*)^ − *ŷ*^(*k*)^∥^2^, where *y*^(*k*)^ ∈ ℝ and *ŷ* ^(*k*)^ ∈ ℝ are the assayed and predicted fitness for variant *k*, respectively. This shift not only demands a change of output space, from the probability distribution space over *X* to the space of all real values R, but also compels the model to reconfigure its learned parameters, often at the cost of erasing its pre-trained features, especially when the volume of fine-tuning data is limited.

Based on this observation, our key idea to address catastrophic forgetting is to align the fine-tuning objective with the pre-training objective. We hypothesize that the pre-trained pLM, already adept at assessing the evolutionary plausibility of sequence mutations, only needs minor adjustments in its output probability distributions to characterize a protein-specific fitness landscape. Therefore, rather than retraining the pLM on a different task (MSE minimization), we maintain the likelihood maximization paradigm and only subtly adjust the pLM’s parameters such that the output probability distribution *p*(*x*_*i*_|*x*_−*i*_) more closely correlate with the given low-*N* fitness data, thereby enhancing the pLM’s prediction accuracy without losing its pre-trained evolutionary patterns.

To gain more intuitions about our method, let us consider a special example of single-mutation variants (Fig 1e). For the *i*-th residue of a wildtype sequence ***x***, a pLM predicts a 20-dimensional probability distribution *p*(*x*_*i*_|*x*_−*i*_). Previous studies ^41;43;51;55^ found that when the wildtype AA *x*_*i*_ mutated to another AA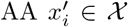, the predicted probability 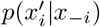(or its normalized version to the wildtype probability) is a reasonable proxy of the mutated sequence’s fitness 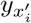. Now, if we are given the fitness scores of some mutations at site *i* as labeled data, we could refine the original distribution *p*(*x*_*i*_|*x*_−*i*_) to become more consistent with the fitness values. For two substitutions 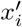 and 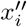 on the same position where 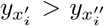,we want the calibrated distribution *p*^*′*^ to preserve the same fitness ordering in the probability space, i.e.,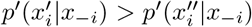. This calibration steers the pre-trained pLM to be more consistent with the fitness data while still respecting its pre-learned evolutionary patterns.

### 3.2 Contrastive Fitness Learning

We now introduce our novel fine-tuning algorithm, dubbed contrastive fine-tuning, to train a pLM-based model for the low-*N* learning of the protein fitness landscape. Formally, in our protein fitness prediction problem, we are given i) a pre-trained pLM 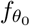 parameterized by *θ*_0_, which predicts the probability *p*_0_(*x*_*i*_|*x*_−*i*_), and ii) a small (low-*N*) set of functionally characterized variants 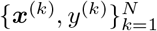 of a particular protein, where ***x***_*k*_’s are the variant sequences and *y*_*k*_’s are the experimentally measured fitness values. Our goal is to calibrate the pre-trained Plm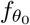, through fine-tuning on the low-*N* fitness data, into an enhanced model *f*_*θ*_ for low-*N* fitness prediction. In this work, we selected ESM-1v ^40^ as the pre-trained pLM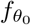 for its effective zero-shot prediction performance in fitness prediction ^40;43^, but other pLMs ^26–28^ can also be used. The neural network architecture of *f*_*θ*_ mirrored that of 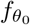, with the parameters *θ* initialized using the pre-trained weights *θ*_0_.

#### From pLM to fitness prediction

Distinct from conventional pLM fine-tuning approaches (e.g., full or top-layer fine-tuning, Figs 1c-d), ConFit maintains the pre-training objective (i.e., predicting probability *p*_*θ*_(*x*_*i*_|*x*_−*i*_)) in fine-tuning, rather than fine-tuning the pLM to directly predict the fitness values *ŷ* using a regression objective. As discussed in Sec. 3.1, this approach circumvents the drastic parameter changes to accommodate the new learning task and output space.

In lieu of directly estimating fitness values, ConFit extracts fitness proxy from the conditional probability *p*_*θ*_(*x*_*i*_|*x*_−*i*_) predicted by pLM *f*_*θ*_. This is informed by prior research ^40;43;51^, which demonstrates the effectiveness of using such probabilities in predicting AA mutation effects. We quantify the impact of a given mutation by comparing its pLM portability with the reference probability of the wild type. Specifically, let ***x***^MT^ and ***x***^WT^ denote the mutant and wild-type sequences, respectively, and *M* the mutated sites. The effect of a substitution 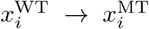 is quantified by the log probability ratio at this site: 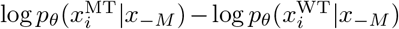. Consequently, we aggregate the mutational effects to approximate the evolutionary plausibility of the mutant sequence:

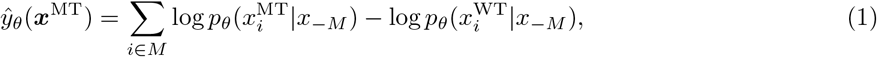

Previous work ^20;37;41;43^ found that this score is an informative proxy of the fitness of mutant ***x***^MT^.

#### Contrastive fine-tuning

To fine-tune *f*_*θ*_ for fitness prediction, instead of re-learning a model to minimize the MSE between predicted and true fitness values, we propose to *calibrate* the pLM probability distribution *p*_*θ*_(*x*_*i*_|*x*_−*M*_), adhering to the original pre-training objective of likelihood maximization. Our calibration strategy is inspired by contrastive learning, an ML strategy that compares multiple outputs of a model and encourages them to be ordered according to some criteria. In our case, we aim for a calibrated model where the likelihood-derived evolutionary scores align with the experimental fitness values: given two samples (***x***^+^, *y*^+^) and (***x***^−^, *y*^−^) with *y*^+^*>y*^−^, we want to have *ŷ* _*θ*_(***x***^+^)*>ŷ* _*θ*_(***x***^−^). This objective can be described by a loss function defined based on a classic probability model known as Bradley–Terry (BT) model ^56^, which we term the BT loss:

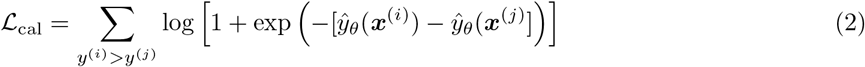

This loss function encourages the model to rank predicted fitness scores in the correct order relative to the ground truth. Although ConFit is not directly trained to predict the precise *magnitude* of fitness values, its prediction of fitness *rankings* is particularly valuable in the context of protein engineering, where ML models are often employed to efficiently rank a set of candidate variants *in silico*, thereby prioritizing the most promising variants for further experimental screening. Moreover, employing pairs from the *N* samples to compute the BT loss increases the effective training size from *O*(*N*) to *O*(*N* ^2^), further alleviating the data scarcity. Several other (scaled) ranking losses ^57^ can be explored in future work in place of ℒ _cal_. A similar

BT loss was also used in recent studies for protein sequence design ^58^ and fitness epistasis learning ^59^.

#### Regularization

To better avoid catastrophic forgetting, we further introduce another Kullback–Leibler (KL) divergence-based regularization loss to prevent the calibrated distribution *p*_*θ*_ deviating too far away from the pre-trained distribution 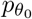 :

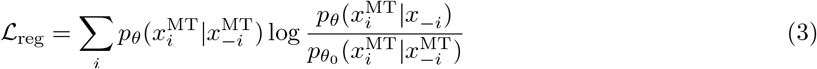

Our final loss function is a linear combination of the calibration loss and the KL regularizer: *ℒ* = *ℒ* _cal_ +*λℒ* _reg_, where *λ* is a coefficient that balances the two losses.

*Remarks*: Our fine-tuning algorithm is partially inspired by a key technique known as Reinforcement Learning with Human Feedback (RLHF) ^60^ that enables ChatGPT to generate high-quality texts. In ChatGPT, human feedback about generation quality (e.g., text A is better than text B) is used to fine-tune GPT to align with human preferences. Similarly, in ConFit, we use experimental fitness data (e.g., mutation A leads to higher fitness than mutation B) to fine-tune a pLM into a low-*N* fitness prediction model.

### 3.3 Efficient Fine-Tuning with Low-Rank Reparameterization

The pre-trained pLM used in ConFit (ESM-1v) has 650 million parameters, and updating all parameters during our contrastive fine-tuning can be computationally expensive. We thus employed Low-Rank Adaptation (LoRA) ^61^, a parameter-efficient fine-tuning (PEFT) method to reduce the number of updated parameters. LoRA factorizes the weight updates to each neural network layer in ESM1v, denoted as a matrix 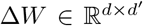, into two low-rank matrices Δ*W* = *UV* where *U* ∈ ℝ^*d×r*^, 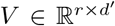, and the rank *r*≪ min(*d, d*^*′*^), thereby reducing the number of trainable parameters from *O*(*d × d*^*′*^) to *O*((*d* + *d*^*′*^) *× r*). In our experiment, we applied LoRA to ESM-1v with rank *r* = 8, reducing the effective number of parameters to 1.35 million, a 99.79% reduction compared to the 650 million parameters in the full ESM-1v model (Supplementary Information).

In addition to its advantages of computational efficiency, the reduction in trainable parameters holds significant implications for ConFit’s adaptation to make protein-specific fitness predictions, especially in low-data scenarios. Recent studies in ML suggested that updates to language model weights often exhibit a low ‘intrinsic rank’ ^61^ and PEFT methods enable superior performance in few-shot learning tasks compared to conventional full-parameter fine-tuning methods ^61–65^. While the vast capacity of the 650 million parameters enables ESM-1v to capture complex patterns in protein fitness landscapes, it could be over-parameterized when dealing with limited fitness data. Fine-tuning the entire model in such cases risks catastrophic forgetting. We thus leveraged LoRA to exploit the low-rank structure of pLM parameters and achieve efficient adaptation to new proteins using low-*N* data.

### 3.4 Enhancing Fitness Prediction with MSA Context Retrieval

While the pre-trained pLM used in ConFit captured global evolutionary patterns across natural protein sequences, when predicting fitness for a protein of interest, we further retrived its evolutionary-related sequences (homology) to reinforce the local evolutionary contexts specific to this protein.

Specifically, starting with the wildtype sequence of the protein, we searched in the UniProt protein sequence database for its homology and built a multiple sequence alignment (MSA). Next, we fit DeepSequence ^20^, a variational autoencoder (VAE)-based density model, on the MSA to estimate sequence likelihood *p*_MSA_(***x***). Different from the pLM sequence likelihood estimated from all natural sequences, the MSA-based sequence likelihood *p*_MSA_ here was only fit on the target protein’s MSA sequences, thus capturing the local evolutionary contexts specific to this protein. Similarly, DeepSequence used the log-odd ratio to predict the fitness of a variant: *ŷ*_MSA_(***x***^MT^) = log *p*_MSA_(***x***^MT^) − log *p*_MSA_(***x***^WT^), which have been shown as competitive unsupervised fitness predictor ^20;22^. A recent study leveraging both pLM and retrieval of MSA sequences achieved state-of-the-art performance on zero-shot protein fitness prediction^42^.

In ConFit, we refined the pLM-based fitness prediction *ŷ*_*θ*_(***x***) by fusing it with the MSA-based prediction *ŷ*_MSA_(***x***): *ŷ* (***x***) = α*ŷ*_*θ*_(***x***) + (1 − *α*) *ŷ*_MSA_(***x***), where *α* is a weighting factor. We empirically determined the value as *α* = 0.8 by inner-loop cross-validation. Note that ConFit integrates multi-scale evolutionary contexts to predict protein fitness, including the global scale (natural protein sequences), intermediate scale (MSA sequences), and the most specific scale (protein-specific fitness data).

### 3.5 Implementation Details

#### ESM Ensemble

The ESM-1v study ^40^ released five sets of pre-trained model weights, all based on the same model architecture but initialized with different random seeds. Following their approach, we average the outputs of the five models to derive the fitness prediction *ŷ*_*θ*_ (Eq. 1) (Supplementary Information).

#### Training details and hyperparameter tuning

We trained ConFit on 4 NVIDIA A40 GPUs using the Adam optimizer and a cosine annealing scheduler, which reduced the learning rate from an initial rate to a minimal value. To mitigate the risk of overfitting, we early-stopped the training if the validation performance was not improved. We tuned the hyperparameters using an inner-loop validation data split within the training set, and the choices of hyperparameters were listed in (Supplementary Information).

## 4 Results

We conducted multiple evaluation experiments to demonstrate ConFit’s low-*N* prediction ability for protein fitness and its utility in navigating protein fitness landscapes for protein engineering applications.

### 4.1 Experimental Settings

#### Datasets

To benchmark protein fitness prediction, we downloaded 34 fitness datasets across 27 proteins generated from deep mutational scanning (DMS) or random mutagenesis studies, following the choice of a prior work ^38^ (Supplementary Information). These datasets were originally curated by the DeepSequence study ^20^ and supplemented by datasets from other mutagenesis studies ^66;67^. The fitness-labeled variant sequences range from single or double mutants to more extensively mutated sequences ^67^.

#### Data split

For each fitness dataset, we withheld 20% of randomly sampled data as test set, with the remaining 80% data used as the training library. To simulate varying degrees of low-*N* scenarios, we created the training set by randomly sampling *N* =48, 96, 168, or 240 samples for the training library. These particular sizes were chosen to reflect the common dimensions of laboratory well plates used in experimental assays. Each training set size was tested across 10 independent trials to ensure reproducibility.

#### Baseline methods

ConFit was compared to 13 leading sequence-based protein fitness prediction methods, including supervised and unsupervised models (Supplementary Information). For supervised methods, we included ‘augmented models’ (Augmented VAE and Augmented EVmutation) proposed by Hsu et al. ^37^ which combine amino acid sequences and evolutionary density scores as input features, and are recognized for their state-of-the-art supervised fitness prediction performances ^37;38^. The comparison also included eUniRep, a model specifically tailored for low-*N* applications ^36^. We further included 10 top-ranked unsupervised models from the ProteinGym benchmark for zero-shot fitness prediction ^42^, covering a wide range of modeling techniques, including language models (ESM-1b ^27^, ESM-1v ^40^, UniRep ^25^, WaveNet ^50^, MSA Transformer ^68^), Potts models (e.g., EVmutation ^9^), latent variable models (e.g., EVE ^22^, DeepSequence ^20^), and hybrid models (e.g., Tranception ^42^, TranceptEVE ^69^. Given our focus on sequence-based predictions, recent fitness models requiring structural data inputs ^32;38^ were not included in our comparison.

### 4.2 Accurate Prediction of Protein Fitness

In our first experiment, we evaluated the accuracy of the fitness prediction of ConFit. We restricted the training set size to include only *N* = 240 variants randomly sampled from the training library. In contrast to prior studies ^34^ where methods were trained on copious amounts of training samples (e.g., 80% of data), here we limited the training data to a mere *N* = 240 variants randomly sampled from the training library (14% data on average per datasets), which simulated a more realistic data-scarce scenario in protein engineering. We first compared ConFit to leading unsupervised methods, including ESM1v, EVE, EVmutation, and TranceptEVE. These methods fit generative sequence density models on general natural sequences or specific homologous sequences to estimate the likelihood of a protein sequence and predict the mutation effects by comparing the log-odds ratio between the mutated and wild-type sequences. We found that, despite the small training set size, ConFit effectively harnessed information from the *N* = 240 samples and improved the fitness prediction accuracy without overfitting (Fig 2a). On average, ConFit achieved a mean Spearman correlation of 0.66 across all datasets, outperforming the unsupervised methods with clear margins on 29/34 datasets. In particular, compared to ESM1v – the non-finetuned counterpart of its base model – ConFit substantially improved the prediction accuracy (median difference of Spearman correlation Δ*ρ* = 0.23; Fig 2b), suggesting the effectiveness of our contrastive fine-tuning algorithm. ConFit also significantly outperformed TranceptEVE (median Δ*ρ* = 0.19; Fig S1), the best predictor as evaluated by the ProteinGym fitness prediction benchmark ^42^, and six additional unsupervised methods (Fig. S2).

**Figure 2:**
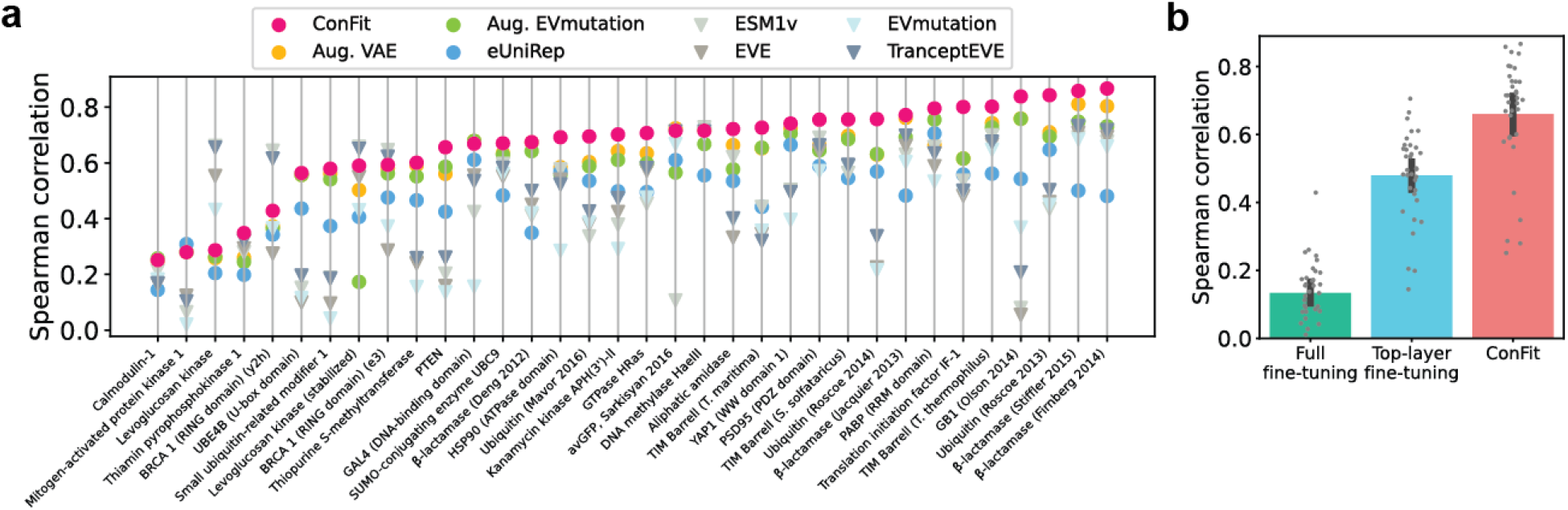
Comparisions between ConFit and other methods for protein fitness prediction. (**a**) Benchmark results on using 34 mutagenesis datasets. Supervised methods were shown as dots and unsupervised methods as triangles. Supervised models were trained on randomly sampled *N* = 240 samples. Each dot represents the average Spearman correlation over 10 repetitions. (**b**) Comparisons between ConFit and conventional fine-tuning strategies. Bar plots represent the mean*±*SD spearman correlations over the 34 fitness datasets.

We then compared ConFit to three supervised methods, including Augmented VAE, Augmented EVmutation, and eUniRep ^36^. Existing studies demonstrated these methods are effective low-*N* fitness predictors ^36;37^. We observed that ConFit was still in a leading position when compared to these supervised baselines, achieving the maximal Spearman scores on 29/34 datasets (Fig 2a). Even on the remaining 5/34 datasets where ConFit was not the top performer, its Spearman scores were very comparable to the top methods, Augmented VAE or Augmented EVmutation (mean Δ*ρ* = −0.01; Fig S1). Moreover, ConFit achieved a 38% increase in Spearman scores compared to eUniRep, a pLM using the top-layer fine-tuning strategy, and a substantial 400% improvement compared to the full fine-tuning strategy (Fig. 2b). This gap affirmed our discussions in Sec. 2: full pLM fine-tuning tends to overfit, and while top-layer fine-tuning mitigates this risk, it can sacrifice predictive precision as it may not fully capture the complex features in protein fitness landscapes. In contrast, ConFit better characterized the fitness landscape with our contrastive fine-tuning strategy, which takes full advantage of pLMs by fine-tuning the entire model while preventing overfitting.

Overall, the improvements over state-of-the-art methods in this evaluation demonstrated the effectiveness of ConFit in adapting pre-trained pLMs for protein fitness prediction.

### 4.3 Low-*N* Learning of Fitness Landscape

Having evaluated the fitness prediction performance of ConFit across 34 DMS datasets in comparison to existing methods, we proceed to specifically assess the low-*N* learning capability of ConFit. We systematically varied the size of the training set by randomly sampling *N* = 48, 96, 168, 240 variants from the non-test data. We first observed that, as expected, ConFit’s prediction accuracy increased with increasing labeled training data (Fig. 3a). Notably, using only *N* = 48 training samples, ConFit predicted protein fitness at a Spearman correlation of 0.56, only 15% lower than its performance at *N* = 240, suggesting ConFit can effectively adapt a general pLM to accurately characterize a protein-specific fitness landscape even using an extreme low-*N* fitness dataset. The results also suggested that the MSA context retrieval consistently enhanced ConFit’s performance across all training data sizes (Fig. 3a).

**Figure 3:**
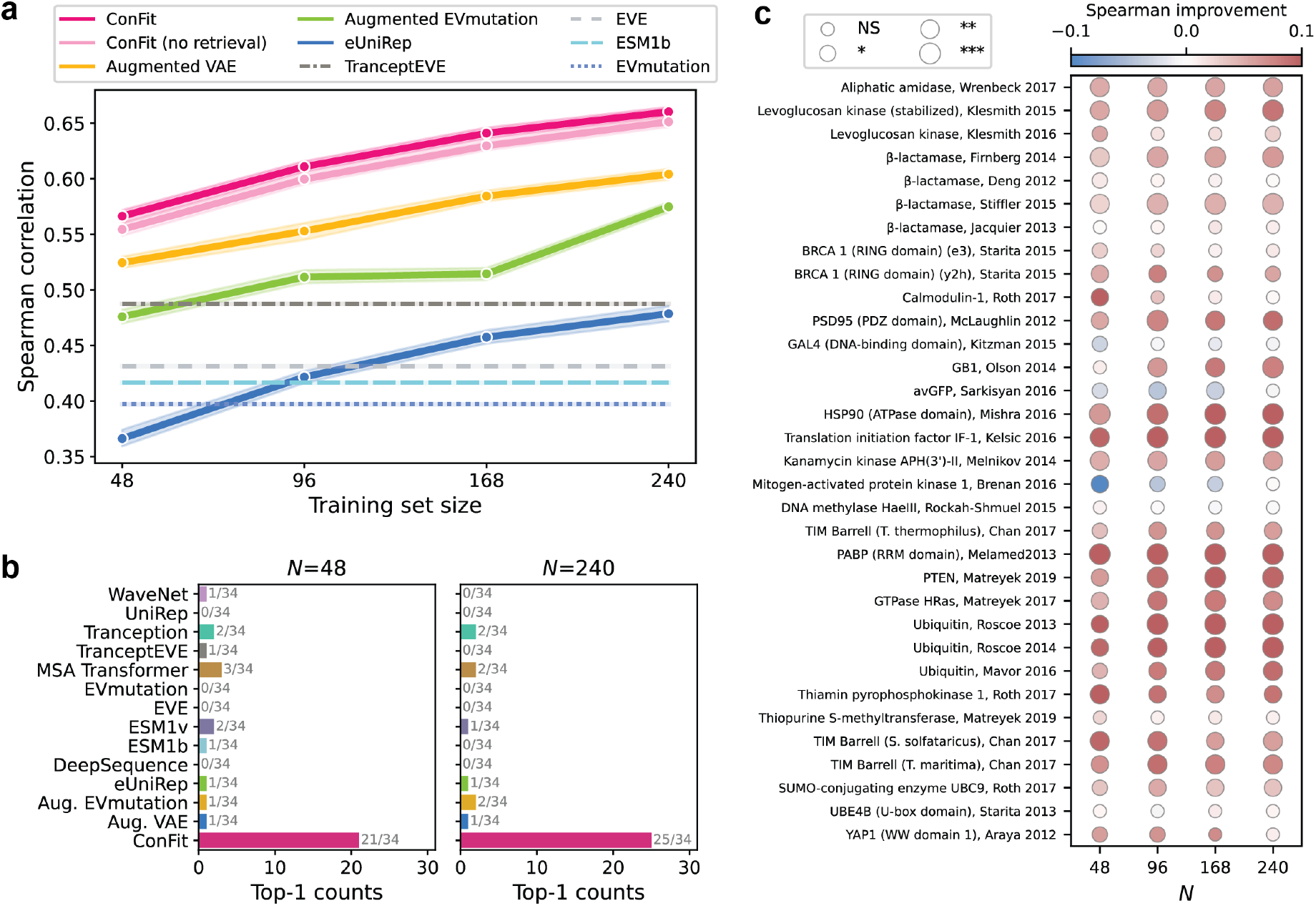
Evaluations on low-*N* learning of protein fitness. (**a**) Prediction performance of methods trained on *N* =48, 96, 168, or 240 samples. Solid lines and error bands indicate the mean*±*SD of Spearman correlations achieved by supervised methods. Performances of unsupervised methods are shown in dashed lines. (**b**) The frequency of each method ranked as the best across 34 datasets for training set sizes 48 and 240. (**c**) Detailed comparison between ConFit and Augmented VAE. The heatmap shows the improvements in Spearman correlation achieved by ConFit for each dataset and each training set size *N* . The dot size indicates the statistical significance. NS: not significant; ^*^: *P* ≤ 0.05; ^**^: *P* ≤ 0.01; ^***^: *P* ≤ 0.001.

Compared to baseline methods, ConFit consistently outperformed three supervised methods and ten unsupervised methods across various training data sizes (Figs. 3a and S3). The performance improvements achieved by ConFit in low-*N* fitness learning were also substantial: for training data sizes *N* = 48, 96, 168, 240, ConFit achieved the top-1 Spearman correlation on 21, 24, 26, and 25 out of the 34 DMS datasets, respectively (Figs. 3b and S4), while second best baseline ranked at the top for only 3/34 datasets. We further analyzed the prediction performances of ConFit and Augmented VAE, the state-of-the-art supervised method for low-*N* fitness prediction ^37^. ConFit outperformed Augmented VAE across all training sizes *N* on 28/34 DMS datasets and outperformed for at least one *N* value for 31/34 DMS datasets, and the majority of the improvements were statistically significant (Fig 3c).

Together, these results demonstrate the superior low-*N* fitness learning ability of ConFit. ConFit’s performance gains stem from our novel contrastive fine-tuning algorithm that prevents overfitting without compromising prediction accuracy. Our approach significantly advances the current paradigm of low-*N* protein fitness prediction. The prevailing strategy employed by prior methods to achieve low-*N* fitness learning is to use informative pLM embeddings or evolutionary scores derived from unsupervised models as input, while keeping the predictive model in the simplest form (e.g., linear regression) ^36;37^. The rationale behind this strategy is that models with fewer parameters are less prone to overfitting on low-*N* data. Although not overfitting, this strategy compromises prediction accuracy due to the limited effectiveness of the predictive models. ConFit, however, completely changed this paradigm through the contrastive finetuning of pLMs – the pLM in ConFit is trainable, rather than being fixed as in previous methods ^36;37^, and carefully tailored through the contrastive fine-tuning algorithm to better modeling the complex sequence-fitness relationships, boosting the prediction accuracy without overfitting.

### 4.4 Extrapolation from Single to Higher-order Mutants

A critical metric for evaluating protein fitness prediction methods is their generalizability from lower-order mutants to higher-order mutants, which is vital for guiding protein engineering to discover novel higherorder mutants based on the fitness data of lower-order mutants. Among the 34 fitness datasets, we selected three—pertaining to avGFP, GB1, and PABP proteins—that included high-order mutants and conducted an evaluation where the model trained on single-mutation data is used to predict fitness for high-order mutants (Fig 4a). To further challenge the evaluated methods, we simulated the low-*N* scenarios with limited training data size *N* = 48, 96, 168, 240. We found that ConFit exhibited an outstanding extrapolation performance, with substantially improved or competitive performance as compared to other supervised baselines (Fig 4b). With increasing *N*, ConFit also increased in prediction accuracy and the performance gap over other methods.

**Figure 4:**
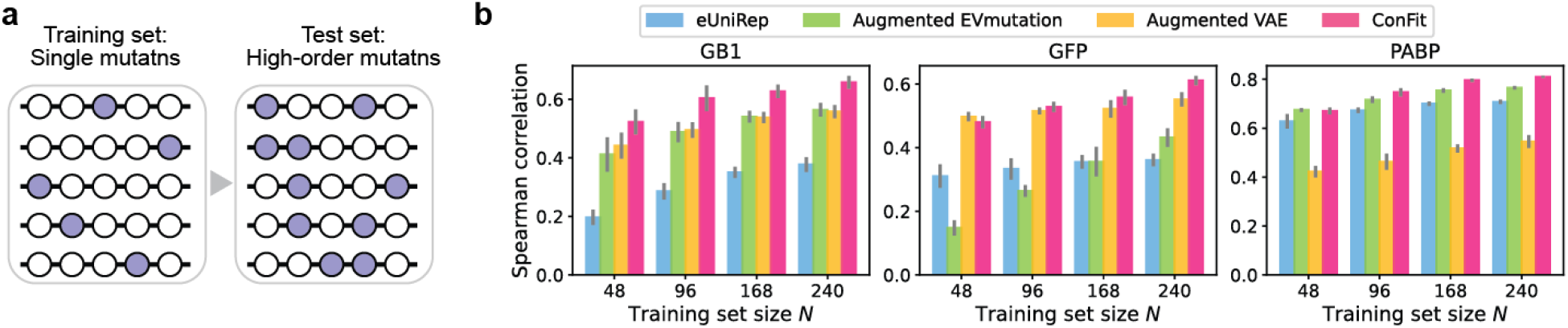
Extrapolation performance of fitness prediction on multi-mutation variants. (**a**) Each method was trained using the fitness data of single mutants data and then tested on higher-order mutants. (**b**) Comparisons between ConFit and three supervised baseline methods on three proteins (GB1, GFP, and PABP). Randomly sampled *N* =48, 96, 168, or 240 single mutants were used as training samples.

The strong extrapolation performance of ConFit is particularly useful for comprehensively characterizing the protein fitness landscapes. Current DMS studies often only sample point-mutation variants or variants within a local sequence space, mainly due to the extensive experimental efforts associated with the generation and analysis of higher-order variants. ConFit thus offers an effective solution to extrapolate from low-order mutants to identify higher-order functional mutants in protein engineering.

### 4.5 Navigating Fitness Landscape with Active Learning

Given the outstanding low-*N* learning capability and extrapolation performance of ConFit, we next demonstrated its utility in protein engineering using an active learning experiment, in which the goal is to strategically select training variants such that better fitness prediction performance can be achieved with less data. Active learning simulates the “prediction–acquisition–screening–retraining” loop in ML-guided protein engineering applications: scientists iterate between ML prediction and experimental screening for multiple rounds; in each round, the ML model proposes promising variants for screening, and the screened fitness data are then utilized to retrain the ML model, which in turn informs the subsequent cycle of predictions.

We evaluate ConFit’s efficiency for landscape exploration on three proteins (GB1, PTEN, and UBE4B) with various definitions of fitness (binding, stability, and Ubiquitin ligase activity, respectively). We again set aside 20% of the data in each dataset as test data, reserving the remaining 80% as a training library from which we can draw samples. We initiated the active learning loop by training ConFit on *N* = 48 randomly sampled variants. In each subsequent round, ConFit predicted the fitness for all test variants and prioritized 48 variants from them, which were then added to the training set with their ground-truth fitness values for retraining ConFit. The prioritization was based on a scoring function called upper confidence bound (UCB), defined as *s*(***x***) = *ŷ* (***x***) + *β σ* (*ŷ* (***x***)), where *ŷ* (***x***) is ConFit’s fitness prediction for sequence ***x***, and *σ* (*ŷ* (***x***)) quantifies the prediction uncertainty using the standard deviation of outputs from the five ESM-1v models within ConFit. The coefficient *β* balances the tradeoff between high predicted fitness (exploitation) and high uncertainty (exploration). We set *β* = 1 in our experiments. For comparisons, we included a greedy scoring function that ranks variants based on the predicted fitness and a random scoring function that randomly samples variants from the training library.

Test results showed an increasing trend in ConFit’s performance through five iterations of active learning on all three proteins (Fig 5). Interestingly, the random scoring function did not yield an increasing trend in performance, especially on GB1 and UBE4B. This indicated that the quality of the training set matters more than its quantity for improving the model’s prediction accuracy. Since the protein sequence space contains a large fraction of non-functional sequences, the random strategy easily sampled non-functional sequences, which are less informative for model training. Additionally, the greedy sampling strategy was consistently outperformed by UCB, suggesting that only exploiting high-fitness regions in the sequence space may restrict the model’s generalizability to unexplored regions. In contrast, using UCB, ConFit prioritized not only variants likely to deliver high fitness but also those that can best address the model’s uncertainty for improving prediction accuracy.

**Figure 5:**
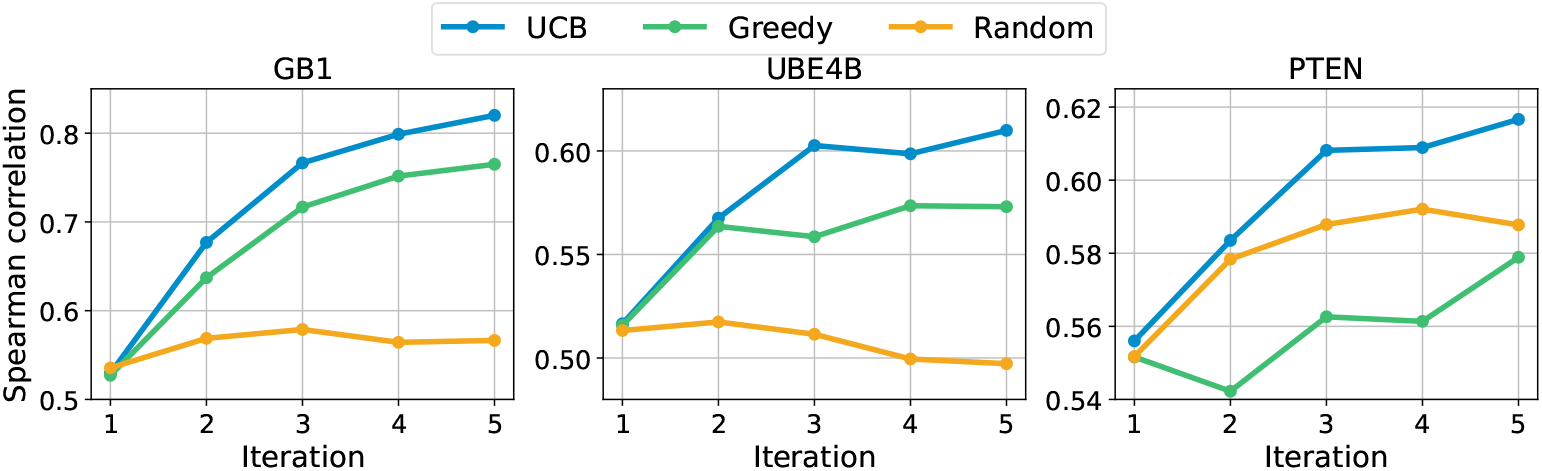
ConFit efficiently characterizes fitness landscapes with active learning. *In silico* active learning experiments were simulated on three protein fitness landscapes. In each iteration, variants prioritized by ConFit and a certain data acquisition function (Upper Confidence Bound, Greedy, or Random) were labeled with ground-truth fitness and used to augment the training set for model re-training.

The results also demonstrated the advantages of iterative data acquisition and model refinement. For example, on the UBE4B protein, ConFit was progressively refined through active learning using an cumulative number of 240 variants (48 per round *×* 5 rounds) and culminated in a Spearman correlation of 0.609 (Fig 5, UBE4B). However, when trained on the same amount of samples in a single training session, ConFit only achieved a Spearman correlation of 0.497 (Fig 5, UBE4B). This suggested that the accurate low-*N* fitness predictions of ConFit enable a feedback mechanism between ML models and experiments, which leads to more efficient navigation of protein fitness landscapes given the same experiment budget.

## 5 Discussion

We presented ConFit, an ML method for low-*N* protein fitness prediction through contrastive fine-tuning of pLMs. Many state-of-the-art ML solutions for protein science problems now rely on pLMs since pLMs capture the implicit evolutionary, structural, and biophysical constraints in protein sequences, thanks to their large model sizes and training data sizes. However, it is challenging to adapt the general pre-trained pLMs to problem-specific predictive models for protein biology tasks because large pLMs typically tend to overfit on small-size task-specific training data. Observing that ML fitness prediction models are often used in protein engineering to rank variants for prioritizing promising ones for screening, we proposed a contrastive fine-tuning algorithm to reprogram a pre-trained pLM for predicting the relative order of variant fitness. This strategy makes the pLM’s pre-training and fine-tuning objectives consistent, avoiding the catastrophic forgetting issues frequently observed in naive fine-tuning approaches. Extensive evaluations suggested that ConFit outperformed over ten existing ML methods for protein fitness prediction and excelled especially in small-data scenarios. We expect ConFit to serve as an efficient ML algorithm for low-*N* protein engineering. Moreover, our contrastive fine-tuning algorithm represents a new paradigm for fine-tuning pLM better than conventional approaches (full or top-layer fine-tuning) and can be extended to other problems in protein informatics.

