## Supplementary Information for "Contrastive Fitness Learning: Reprogramming Protein Language Models for Low-*N* Learning of Protein Fitness Landscape"

### A Supplementary Information

#### A.1 Datasets

In our study, we examined 34 benchmark deep DMS datasets across 27 proteins, following the settings outlined in a previous study[38]. Among the 34 DMS datasets, the GB1 dataset and the avGFP dataset were collected from previous studies[66;67], while the remaining 32 datasets were curated by the DeepSequence[20]. The GB1 dataset comprises 2,457 mutations, encompassing both single and double mutations exhibiting high epistasis as found in previous studies[38;70]. In addition, the avGFP dataset includes mutations located within positions with more than 30% gaps in MSA, ensuring a focus on regions abundant with evolutionary data[36–38]. Positions 15–150 were selected out of 237 positions, yielding 7,775 mutants with mutation counts ranging from one to nine.

For each dataset, 20% of the labeled data was randomly sampled as the test set. Training sets with various sizes were sampled from the remaining 80% of the data in each run, simulating low-$N$ scenarios in protein engineering. We chose a set of fixed training data sizes ranging from 48 to 240 samples (e.g. 48, 96, 168, 240), consistent with the sizes of commonly used well plates in the laboratory. To ensure the reproducibility of our results, we performed 10 independent repetitions of model training and evaluation for each fixed training data size.

#### A.2 Efficient Fine-tuning with LoRA

The pre-trained parameters of a neural network layer in ESM1v can be described by a weight matrix $W_0 \in \mathbb{R}^{d\times d'}$, where $d$ and $d'$ are the input and output dimensions of this layer, respectively. During the fine-tuning, the update on $W_0$ can be represented as $W_0' = W_0 + \Delta W$, where $\Delta W \in \mathbb{R}^{d\times d'}$ is the update matrix. It has been observed that the weight matrices in a pre-trained LLM often have a low intrinsic rank[71]. Therefore, LoRA[61] constrains the update matrix $\Delta W_0$ to a low-rank matrix: $W_0' = W_0 + \Delta W = W_0 + UV$, where $U \in \mathbb{R}^{d\times r}$, $V \in \mathbb{R}^{r\times d'}$, and the rank $r \ll \min(d, d')$. During the fine-tuning, only the weights in $U$ and $V$ are updated while $W_0$ is frozen, which reduces the trainable parameters to $O((d + d') \times r)$ from $O(d \times d')$. In our experiment, we set the rank $r = 8$ for all layers in ESM-1v and observed that this reduced the effective number of parameters being updated in our fine-tuning to 1.35 million, which is a 99.79% reduction compared to the 650 million parameters in the full ESM-1v model.

#### A.3 Training Details

The ESM-1v study[40] released five checkpoints of the pre-trained models, each initialized with a different random seed. We thus used an ensemble approach to build ConFit: we fine-tuned each of the ESM-1v models using a unique combination of four splits as the training set, rotating one split as the validation set for each model. During inference, the outputs of these five models were averaged to generate the final predictions, enhancing ConFit's prediction accuracy and robustness.

Throughout the training phase, we employed the Adam optimizer and a cosine annealing scheduler, which reduced the learning rate from an initial rate to a minimal value. To mitigate the risk of overfitting, we implemented an early stopping strategy. The hyperparameter values used in our training can be found in the Table S1.

**Table S1:** Hyperparameters used in training. lr: learning rate.

| Training Size | Batch size | Max enduring time | Initial lr | Min lr | Max epoch | $\lambda$ |
|---|---|---|---|---|---|---|
| 48 | 16 | 5 | 5e-04 | 1e-04 | 30 | 0.1 |
| 96 | 16 | 3 | 5e-04 | 1e-06 | 50 | 0.2 |
| 168 | 16 | 3 | 5e-04 | 1e-06 | 55 | 0.2 |
| 240 | 16 | 3 | 5e-04 | 5e-06 | 45 | 0.2 |

#### A.4 Metrics

We used the Spearman correlation coefficient as our primary evaluation metric, which was widely used for evaluating ML methods for fitness prediction in protein engineering[20;40;42;68]. This metric, ranging from −1 to +1, assesses the monotonic relationships between variables. A high Spearman correlation nearing +1 indicated a close alignment in the ranking between the ground truth and predicted fitness, while a low correlation approaching −1 signified dissimilar rankings. The Spearman correlation between two variables, denoted as $X$ and $Y$, is mathematically defined as:

$$\rho(X, Y) = \frac{\text{cov}(R(X), R(Y))}{\sigma_{R(X)}\sigma_{R(Y)}}, \tag{S1}$$

where $R(\cdot)$ denotes the ranking function applied to the variable, $\text{cov}(X, Y)$ represents the covariance between the rank variables, and $\sigma$ are the standard deviation.

#### A.5 Baseline Methods

**EVmutation**: EVmutation[9] is an unsupervised sequence density model that explicitly models the covariations between every pair of residues within a protein sequence. EVmutation employs a pairwise undirected graphical model (Potts model), fitted to the MSA including all homologous sequences associated with the protein of interest. It predicts the impact of single or high-order substitution mutations as the log ratio of sequence probabilities between the mutant and wild-type sequences. In this work, we used the EVmutation implementation provided in the 'plmc' package (May 2018, `https://github.com/debbiemarkslab/plmc`).

**Deepsequence VAE**: DeepSequence[20] utilizes a non-linear Variational Autoencoder (VAE) to model sequence distributions within protein families. Generalizing EVmutation, DeepSequence incorporates a multivariate Gaussian latent variable $z$ to model high-order dependency among residues in a protein and defines the likelihood of a sequence $s$ as:

$$P(s|\theta) = \int P(s|z, \theta)P(z)dz \tag{S2}$$

In practice, DeepSequence approximates the above intractable exact likelihood using the evidence lower bound (ELBO) derived from the VAE[20]. We downloaded the ELBO from the data pre-computed using DeepSequence in a previous work[38].

**Augmented VAE and Augmented EVmutation**: Hsu et al.[37] developed a series of 'augmented' models for fitness prediction. The idea is to fit a regression model on the one-hot-encoded sequence features, augmented with an evolutionary plausibility score derived from a sequence density model. Specifically, given a sequence density model with sequence probability distribution $\log P(s; \varphi)$ parameterized by $\varphi$, the augmented model concatenates the log probability with one-hot site-specific amino acid encodingand fits the following regression model:

$$f(s; \varphi, \theta, \beta) = \beta \log P(s; \varphi) + \theta_0 + \sum_{i=1}^{L} \theta_i(s_i) \tag{S3}$$

where the parameters $\theta_i \in \mathbb{R}^{|\mathcal{A}|}$ , $\theta_0 \in \mathbb{R}$ and $\beta \in \mathbb{R}$ are learned from assay-labeled data. As is shown in their analyses[37], the Augmented VAE and Augmented EVmutation models achieved the best performance in protein fitness prediction. We thus chose those two models as our baseline methods.

**eUniRep**: eUniRep[36] is a regression model for low-$N$ protein engineering using sequence embeddings generated by a PLM known as UniRep[25]. UniRep first trains an LSTM-based PLM on protein sequences from the UniRef50 database. Subsequently, this model undergoes a process known as 'evotuning', wherein it is fine-tuned using homologous sequences of a specific protein of interest. This fine-tuning step refines the model's representation learning. Utilizing this refined model, eUniRep uses the vector representations generated by UniRep for individual protein sequences, together with their fitness values, as the training data to fit a ridge or LASSO regression model. Since the regression model only contains a small number of parameters to learn, this approach does not suffer from severe overfitting in low-$N$ scenarios. In this work, we obtained the evo-tuned model weights of UniRep from [38] and further fine-tuned it through ridge regression on DMS datasets. Our fine-tuning strategy followed the protocols outlined in several previous studies for building eUniRep[36–38].

**Unsupervised fitness prediction models**: In this study, we evaluated the performance of 10 state-of-the-art unsupervised fitness prediction models, which were ranked among the top performers on the ProteinGym benchmark for zero-shot fitness prediction[42]. These models represent a diverse array of modeling strategies, including advanced language models (ESM-1b[27], ESM-1v[40], UniRep[25], WaveNet[50], and MSA Transformer[68]), Potts models (EVmutation[9]), latent variable models (EVE[22], DeepSequence[20]), as well as hybrid models that combine Potts models and latent variable models (Tranception[42] and TranceptEVE[69]). We obtained the predicted scores of these models from the ProteinGym benchmark.

### B Supplementary Figures

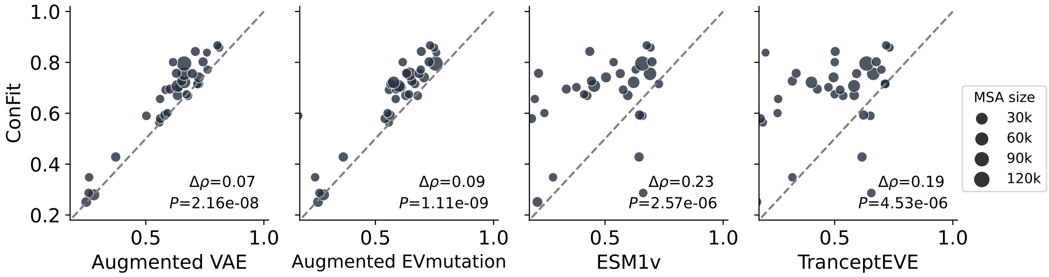


**Figure S1: Pairwise comparison between ConFit and leading supervised and unsupervised methods.** Each point represents the performance on a mutagenesis dataset and the dot size is proportional to the number of homologous sequences in the protein's multiple sequence alignment (MSA). One-sided rank-sum test was used to test the statistical significance.

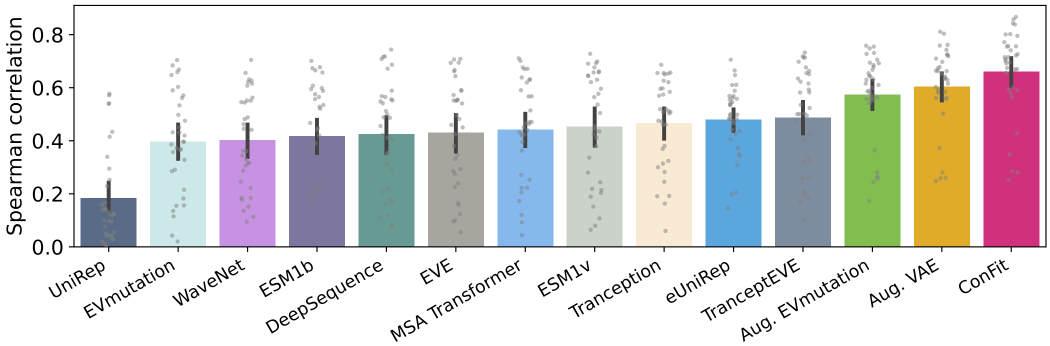


**Figure S2: Full benchmark results for protein fitness prediction.** Supervised models were trained on randomly sampled $N = 240$ samples with 10 repetitions. Barplots and error bars show the mean±SD of Spearman correlations over the 34 fitness datasets.

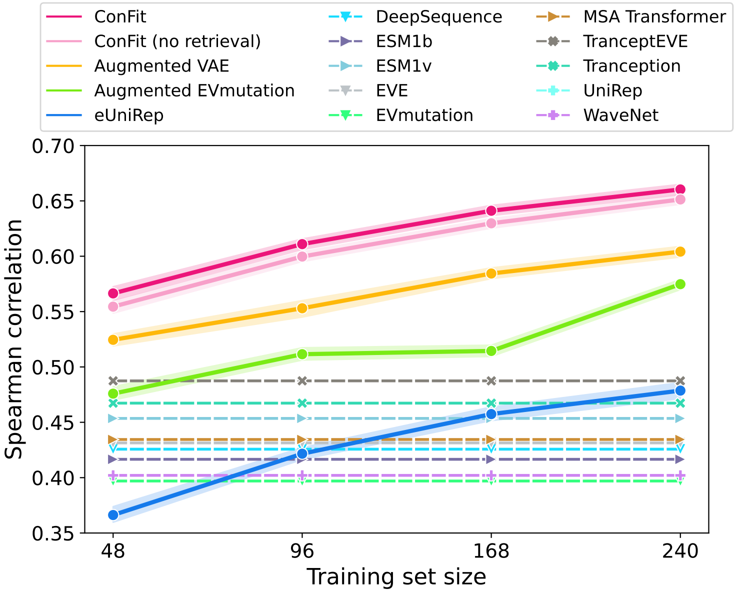


**Figure S3: Low-$N$ prediction performance of all methods.** Supervised models were trained on $N$ =48, 96, 168, or 240 samples. Solid lines and error bands indicate the mean±SD of Spearman correlations achieved by supervised methods. Performances of unsupervised methods are shown in dashed lines.

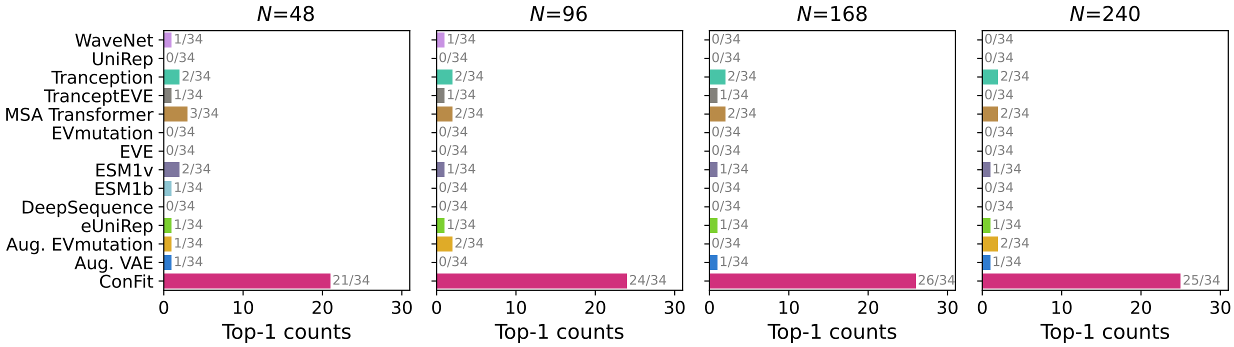


**Figure S4: Frequency of each method ranked as the best**. The frequency of each method ranked as the best in terms of Spearman correlation across 34 datasets for training set sizes 48, 96, 168, and 240
